# Differential regulation of translational stress responses by herpesvirus ubiquitin deconjugases

**DOI:** 10.1101/2025.03.04.641470

**Authors:** Jiangnan Liu, Carlos Ayala-Torres, Maria G. Masucci

## Abstract

The strategies adopted by viruses to counteract the potential antiviral effects of the ribosomal quality control (RQC) that regulates the fidelity of protein translation, ribosome recycling, and the activation of ribosomal and integrates stress responses are poorly understood. Here, we investigated the capacity of the viral ubiquitin deconjugase (vDUB) encoded in the large tegument protein of human pathogenic herpesviruses to interfere with the triggering of RQC upon the induction of translational stress in cytosolic and endoplasmic reticulum (ER)- associated ribosome. We found that the capacity of the vDUBs encoded by Epstein-Barr virus (EBV), Human cytomegalovirus (HCMV), and Kaposi sarcoma virus (KSHV) to counteract the ubiquitination of RPS10, RPS20, and RPS3, and the UFMylation of RPL26 in cells treated with translation elongation inhibitor anisomycin (ANS) correlated with the rescue of model RQC and ER-RQC substrates from proteasome- and lysosome-dependent degradation, promoted the readthrough of stall-inducing mRNAs, and prevented ER-phagy. In contrast, while rescuing the RQC substrate and inhibiting the ubiquitination of RPS10, RPS20, and RPS3 as almost efficiently as the homologs, the Herpes simplex virus-1 (HSV1) encoded vDUB failed to counteract RPL26 UFMylation and did not rescue the ER-RQC substrate or inhibit ER-phagy. Thus, it appears that the different lifestyles of herpesviruses may have promoted the development of distinct strategies for coping with the antiviral effect of ribosome stress responses.

## Introduction

The eukaryotic cell has developed sophisticated mechanisms to monitor and maintain the proteome in physiological and stress conditions via the coordinated regulation of protein synthesis and degradation (1). During viral infection, proteostasis is severely challenged by the antagonistic need of viruses to reprogram the translation machinery towards the rapid production of large amounts of viral proteins and the cellular attempt to curb infection through activation of the PKR kinase that upon the detection of viral nucleic acids, phosphorylates the translation initiation factor eIF2α and represses translation (2). The capacity of viruses to simultaneously induce a profound decline of host protein synthesis via selective shutoff (3) and inhibit PKR (4) highlights the need to maintain high levels of active ribosomes capable of sustaining the translation of viral mRNAs. Although a remarkable array of viral tactics for rapidly subverting translation initiation, elongation, and termination has been described (5), very little is known about the molecular events and viral effectors that may be involved in the regulation of translational stress responses.

The translation of viral mRNAs is challenging due to the presence of long repetitive sequences, complex secondary structures, suboptimal codon usage, and the frequent occurrence of frameshifting and nucleotide misincorporations that can slow down translation or induce ribosome stalling (6). While pausing the elongation cycle may resolve minor issues, persistent stalling may trigger ribosome stress response (RSR) that, via activation of the JNK and p38 kinases, elicit inflammation and may ultimately cause cell death (7). The cells have developed molecular sensors of ribosomal stress and rescue strategies that, via activation of the ribosome-associated quality control (RQC), split the stalled ribosomes, promote the disposal of both damaged mRNA and aberrant translation products, and allow the recycling of ribosome subunits (8, 9). The RQC is initiated by the recognition of collided di- and trisomes (10) or arrested ribosomes (11) or by ligases that perform site-specific mono-ubiquitination of the 40S particle (12–16). In mammals, the RING domain ligase ZNF598 ubiquitinates the RPS10 (eS10) and RPS20 (uS10) proteins (12, 13, 17–19), which elicits splitting of the ribosome subunits by the ASC-1 complex (20, 21), recognition of the 60S-peptidyl-tRNA conjugate by NEMF for CAT tailing (22), ubiquitination of the nascent peptide by the Listerin ligase (23), its extraction form the conjugate by the VCP/p97 chaperone (24), and degradation by the proteasome (8, 25). Following activation of the ISR (26) or when ribosomes are arrested at the initiation codon (16), the RNF10 ligase ubiquitinates RPS2 (uS5) and RPS3 (uS3), which targets the 40S particle for proteasomal degradation in a process known as initiation RQC (iRQC) that is inhibited by the USP10 ubiquitin deconjugase (16, 27).

The stalling of ribosomes that synthesize proteins co-translationally inserted into the endoplasmic reticulum (ER) activates a different branch of RQC, named ER-RQC, dependent on the post-translational modification of the 60S subunits by the ubiquitin-like modifier-1 (UFM1) (28, 29). Although the components, recognition signals, and detailed organization of the ER-RQC are still poorly understood, the UFMylation of RPL26 (uL24) by the UFL1/DDRGK1/CDK5RAP3 ligase complex (30–33) was shown to initiate a signaling cascade leading to either extraction of the nascent polypeptide to the cytosol for proteasomal degradation (34) or to lysosomal degradation of the peptide alone (35) or together with a small portion of the ER in a process known as ER-phagy (36, 37). Failure of the latter pathway following inactivation of the ligase complex by ablation of UFL1 (38, 39), the ER-bound ligase subunit DDRGK1 (30, 35, 40), or the ER-phagy receptor CDK5RAP3 (also known as C53/LZAP) (32, 41), caused the accumulation of immature polypeptides in the ER and initiated the unfolded protein response (UPR) and triggered the ISR. It has been proposed that activation of the ISR in response to failure of the RQC or ER-RQC may counteract the pro-apoptotic effects of RSR by reprogramming the translation machinery in favor of mRNAs containing internal ribosomal entry site (IRES)- or upstream open reading frame (uORF)-dependent that often encode products with pro-survival and anti-apoptotic activity (41–43). It is noteworthy that, although potentially deadly, failure to activate the RQC can, in some cases, be beneficial for the cell because the restart of translation after prolonged pausing facilitates the readthrough of stall-inducing mRNAs (19).

The challenging structure of viral mRNA and the depletion of tRNAs induced by the mass-production of comparatively small proteomes (44) suggest that virus-infected cells may be exquisitely dependent on finetuning of the RQC to secure levels of functional ribosomes sufficient for maintaining efficient virus production. We have previously shown that the viral deubiquitinase (vDUB) encoded in the N-terminal domain of Epstein-Barr virus (EBV) large tegument protein BPLF1 inhibits both RQC (45) and the ER-RQC (46) by regulating the ubiquitination and UFMylation of stalled ribosome, which promotes the translation of viral proteins and enhances the release of infectious virus particles. EBV, also known as human herpesvirus-4 (HHV4), is a lymphotropic virus that establishes latent infections in most human adults worldwide and participates in the pathogenesis of a broad spectrum of lymphoid and epithelial cell malignancies (47) and autoimmune diseases (48). Despite poor genome sequence conservation and significant differences in host ranges and virus life cycles, the large tegument protein of all animal and human herpesviruses investigated encode a functional vDUB (49). Here, we set out to examine whether the vDUBs encoded by EBV, the human cytomegalovirus (HCMV, HHV5), Kaposi sarcoma virus (KSHV, HHV8), and herpes simplex virus (HSV, HHV1) share the capacity to regulate RQC and ER-RQC ribosomal stress responses by interfering with the ubiquitination and UFMylation of ribosomal proteins. We found that the catalytic domain of EBV BPLF1, HCMV UL48, and KSHV Orf64 inhibited the ubiquitination of RPS10, RPS20, RPS3, and RPL26 UFMylation triggered by translational stress, rescued RQC and ER-RQC substrates and impaired ER-phagy. In contrast, HSV1 UL36 selectively failed to prevent RPL26 UFMylation and did not rescue ER-RQC substrates or inhibit ER-phagy, pointing to important differences in the strategies adopted by these human viruses for regulating translational stress responses.

## Materials and Methods

### Reagents

For a complete list of commercial cell lines, reagents, kits, primers, and commercially available or donated plasmids with source identifiers, see Table S1.

### Cell culture and transfection

The HEK293T and HCT116 cell lines were cultured in Dulbecco’s minimal essential medium (DMEM) supplemented with 10% FBS and 10 μg/ml ciprofloxacin (complete medium) and grown in a 37°C incubator with 5% CO2. The cells were transfected using the jetOPTIMUS® DNA transfection reagents according to the protocols recommended by the manufacturers.

### Immunoblotting and co-immunoprecipitation

The cells were lysed for 30 min on ice in NP-40 buffer (50 mM Tris, 150 mM NaCl, 5 mM MgCl2, 1 mM EDTA, 1% Igepal CA-630, 5% glycerol) supplemented with protease inhibitor cocktail. To detect UFMylation, the lysis buffer was supplemented with 10 mM NEM. After centrifugation at 20000xg for 15 min at 4°C, the protein concentration of the supernatants was measured with a protein assay kit. Equal amounts of lysates were fractionated in acrylamide Bis-Tris 4-12% gradient gel (Life Technologies Corporation, Carlsbad, USA). After transfer to PVDF membranes (Millipore Corporation, Billerica, MA, USA), the blots were blocked in Tris-buffered saline (TBS) containing 0.1% Tween-20 and 5% non-fat milk. The membranes were then incubated with the primary antibodies diluted in blocking buffer for 1 h at room temperature or overnight at 4°C, followed by washing and incubation for 1 h with the appropriate horseradish peroxidase-conjugated secondary antibodies. The immunocomplexes were visualized by enhanced chemiluminescence. For immunoprecipitation, the cells were harvested 24 h after transfection and lysed in IP lysis buffer (20 mM Tris-HCl pH 7.6, 150 mM NaCl, 1 mM EDTA, 1% Igepal CA-630, 1% Triton X-100) supplemented with protease inhibitor cocktail for 30 min on ice. For immunoprecipitations under denaturing conditions, the cells were lysed for 10 min on ice in a small volume of lysis buffer supplemented with 1% SDS, followed by dilution to 0,1% SDS. The lysates were incubated with 50 μl anti-FLAG or anti-S Tag conjugates agarose affinity gel at 4°C for 3 h with rotation. The beads were then washed with lysis buffer, and the immunocomplexes were eluted by boiling in 2xNuPAGE Loading buffer supplemented with a sample-reducing agent. All images were acquired using a ChemiDoc Imaging system (Bio-Rad), and the intensity of target bands was quantified using the ImageLab software.

### Ub-VS labeling assay

HEK293T cells were transfected with the vDUBs expressing plasmids in 6-well plates and harvested after 24 h by lysis in 100 µL NP-40 lysis buffer supplemented with a complete protease inhibitor cocktail. Ten µg of cell lysates were incubated with or without 1 µM HA-Ub-VS probe in reaction buffer (50 mM Tris-HCl pH 7.4, 5 mM MgCl2, 1 mM DTT, 0.5% sucrose) for the indicated time at 37 °C. The reaction was stopped by adding 4X NuPAGE sample buffer followed by fractionation in NuPAGE 4-12% gradient gel, as described above.

### Translation-readthrough assay

HEK293T cells were plated into 6-well plates the day before transfection. Semiconfluent monolayers were cotransfected with the FLAG-ev/BPLF1/UL36/UL48/Orf64 and pmGFP-P2A-K0-P2A-RFP or pmGFP-P2A-K20-P2A-RFP and cultured 24 h before detection of GFP and RFP fluorescence by flow cytometry using a BD LSR II SORP apparatus. The data were analyzed using the FlowJo software.

### ER and ER-RQC reporter assay and Endo-H treatment

The RQC K20 reporter was adapted from the ER-K20 reporter by removing the signal peptide and glycosylation site sequences using the Q5® Site-Directed Mutagenesis Kit according to the recommended protocol. The K20 or ER-K20 reporters were cotransfected in HEK293T cells with either FLAG-Tagged vDUBs or the empty FLAG-vector as control. Where indicated, 100 nM Bafilomycin-A1 (BafA1) or 100 nM Carfilzomib were added to the cultures overnight to inhibit lysosome- or proteasome-dependent degradation, respectively. To assess peptide glycosylation, aliquots of the cell lysates were treated with Endoglycosidase H (Endo H) (New Englands BioLabs, Cat#P0702S) according to the manufacturer’s recommendation. Briefly, 10 μg of total cell lysates were denatured at 100 °C for 10 min in the kit’s glycoprotein denaturing buffer. The denatured lysates were treated with 500 units of Endo H at 37 °C for 1 h. The reactions were stopped by adding NuPAGE loading buffer supplemented with a reducing agent and boiling. Samples were analyzed using NuPAGE 4-12% gradient gel electrophoresis.

### ER Autophagy Tandem Reporter (EATR) assay

The expression of the EATR reporter was induced in HCT116-EATR cells (46) grown on cover slides by treatment for 24 h with 2 μg/ml Doxycycline before FLAG-Tagged vDUBs transfection. After 8 hours of transfection, the cell culture medium was removed by repeated PBS washing, and the cells were starved by culture for 16 h in Earl’s balanced salt solution (EBSS) medium with or without Bafilomycin A1. The cells were then fixed and stained with the anti-FLAG antibody as described. Images were acquired using a Zeiss LSM900 confocal microscope, and the number of red dots in 45 vDUBs positive or negative cells was counted manually.

### Statistical analysis

Plotting and statistical tests were performed with data obtained in two or more independent experiments using the Microsoft Excel software or GraphPad Prism 10. No assumptions about data normality were made, and an unpaired two-tailed Student t-test was used to determine statistical significance. Statistical significance is indicated in figures as ns *P* > 0.05, \**P* ≤0.05, \*\**P* ≤ 0.01, \*\*\**P* ≤0.001.

## Results

### The vDUBSs inhibit the RQC and promote the readthrough of stall-inducing mRNAs

To investigate the capacity of the herpesviruses encoded vDUBs to interfere with the RQC responses, we used a pair of reporter plasmids encoding in frame the green fluorescent protein (GFP) and cherry fluorescent protein (ChFP) separated by a linker containing twenty AAA encoded Lys residues (K20). Progression of the translation ribosome through the poly(A) stretch causes ribosome stalling, which, following collision with the trailing ribosomes, activates the RQC and promotes the degradation of the stalled translation product by the proteasome (12). To distinguish between cytosolic and ER-associated events, the reporters expressed the GFP-K20-ChFP polypeptide alone (hereafter K20, Figure 1A) or preceded by an ER-localization signal and glycosylation site (hereafter ER-K20, Figure 3A), which selectively promotes the stalling ribosomes that translate ER-targeted proteins. Previously described FLAG-tagged versions of the relatively well-conserved catalytic domains of EBV-BPLF1 (aa 1-235), HCMV UL48 (aa 1-264), KSHV Orf64 (aa 1-205), and HSV1 UL36 (aa 1-287) (45) were used to assay the effect of the vDUBs on the RQC induced by treatment with a concentration of the translation inhibitor anisomycin (ANS) that promotes random ribosome stalling and collision (18).

**Figure 1.**
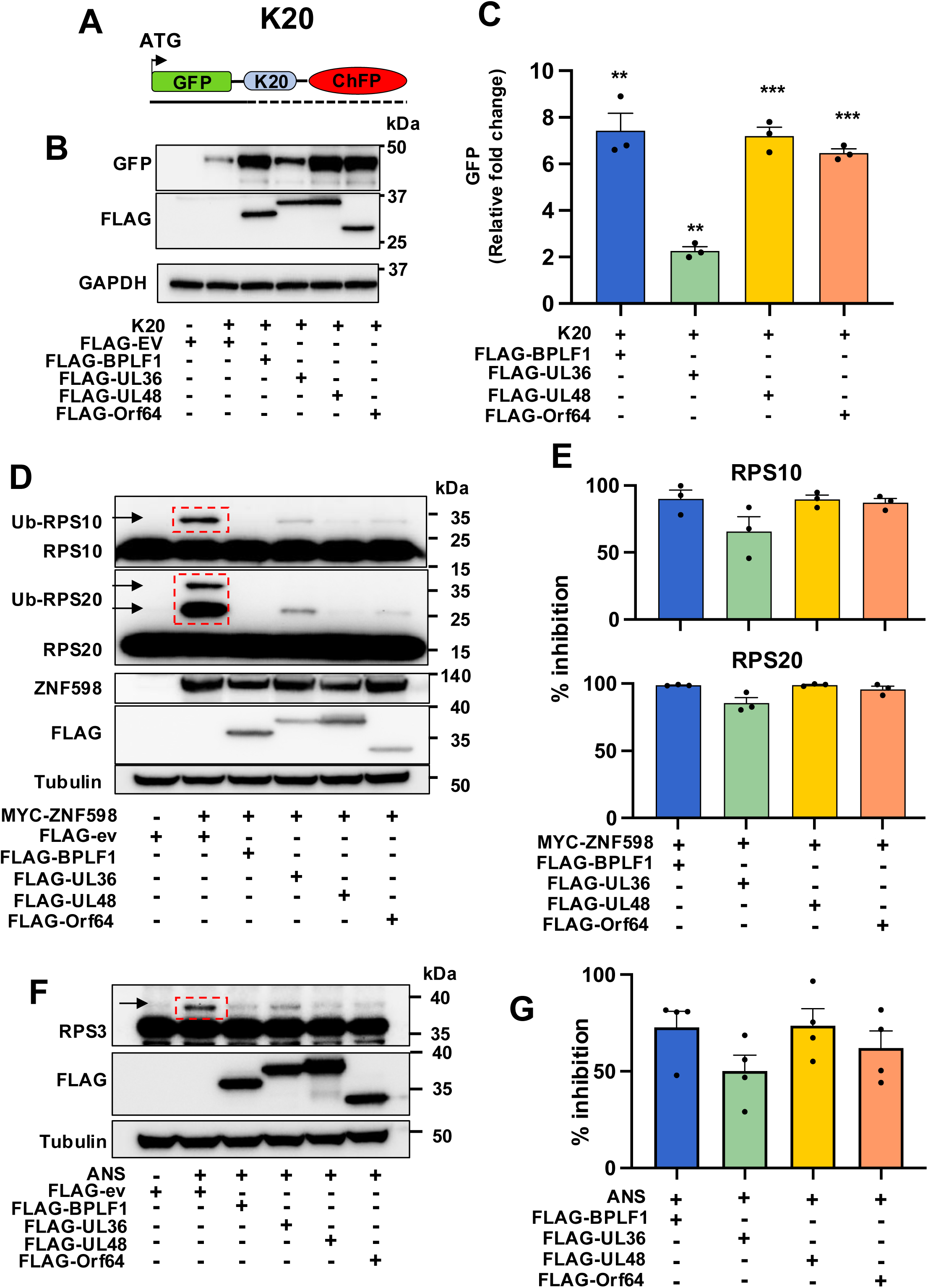
Herpesvirus deconjugases inhibit the RQC. (**A**) Schematic illustration of the RQC reporter (K20). The reporter expresses in-frame an N-terminal GFP following a stretch of Lys residues encoded by AAA codons (K20) and ChFP. Stalling of the ribosome at the poly(A) sequence traps the nascent polypeptide during translation elongation. (**B**) HEK293T cells were co-transfected with the K20 and FLAG-ev/vDUBs plasmids for 24 h, and an equal amount of lysates were analyzed in western blot and probed with indicated antibodies. Blots from one representative experiment out of three are shown in the figure. (**C**) The Mean ± SE relative intensity of the GFP bands in FLAG-vDUBs transfected cells versus in FLAG-ev transfected cells after normalized to loading control in three independent experiments is shown. Significance was calculated by unpaired, two-tailed Student t-test. (**D**) The vDUBs inhibit the ubiquitination of 40 S ribosome proteins by ZNF598. *ZNF598* knockout HEK293T cells were transfected with a ZNF598 expressing plasmid together with FLAG-ev/vDUBs. After 24 h, the cells were lysed in a buffer containing NEM and iodoacetamide to inhibit DUB activity, and western blots were probed with the indicated antibodies. Blots from one representative experiment out of three are shown in the figure. **(E)** The intensities of the Ub-RPS10 and Ub-RPS20 bands (red dotted box in Figure 1D) were quantified by densitometry in three independent experiments. The mean ± SE % inhibition in vDUBs transfected versus FLAG-ev transfected cells is shown. (**F**) The vDUBs inhibit the ubiquitination of RPS3. HEK293T cells transfected with FLAG-ev/vDUBs were treated with 50 ng/ml ANS for 30 min, followed by lysis in a buffer containing NEM to inhibit DUB activity. Western blots were probed with the indicated antibodies. Blots from one representative experiment out of four are shown in the figure. (**G**) The intensities of the Ub-RPS310 bands (red dotted box in Figure 1F) were quantified by densitometry in four independent experiments. The mean ± SE % inhibition in vDUBs transfected versus FLAG-ev transfected cells is shown.

In agreement with our previous finding that BPLF1 and the homologs could rescue from proteasomal degradation an RQC-reporter expressing a GFP construct lacking the stop codon (45), cotransfection of the K20 reporter with FLAG-tagged BPLF1, UL36, UL48 or Orf64 resulted in significant stabilization of a polypeptide of size only slightly larger than full-length GFP (Figure 1A,1B) that is presumably derived from translational arrest at the poly(A) sequence. During translation elongation, the RQC is triggered by the recognition of stalled and collided ribosomes by the ZNF598 ubiquitin ligases that ubiquitinates specific sites on the 40S proteins RPS10 and RPS20 (19). To assess whether the stabilization of the reported correlated with the capacity of the vDUBs to interfere with these ZNF598-dependent ubiquitination events, the expression of ZNF598 was reconstituted in a previously described ZNF598-KO cell line (45) by cotransfection with a ZNF598 expressing plasmid and plasmids expressing the FLAG-tagged vDUBs or, as control, the empty FLAG vector (FLAG-ev). As expected, overexpression of the reconstituted ZN598 resulted in significant ubiquitination of RPS10 and polyubiquitination of RPS20 in cells cotransfected with FLAG-ev, which was almost entirely reversed by co-expression of the vDUBs (Figure 1C, 1D), supporting the conclusion that the viral enzymes share the capacity to counteract the activity of this RQC ligase. The RNF10-dependent ubiquitination of RPS3 was investigated in HEK293T cells transfected with FLAG-ev/BPLF1/UL36/UL48/Orf64 following induction of translation arrest by treatment with 50 ng/ml ANS for 30 min. A comparable inhibition of RPS3 ubiquitination was observed in the vDUB-expressing cells (Figure 1E and 1F). In line with the effect on ZNF598-mediated ubiquitination observed in reconstituted ZNF598-KO cells, the vDUBs inhibited the ubiquitination of RPS10 (Figure S1). However, the effect was somewhat weaker, likely due to the presence of a significant proportion of non-transfected cells. It is noteworthy that although the efficiency of the HSV1 encoded vDUBs appeared to be weaker in both the K20 stabilization and ribosome protein deubiquitination assays, the differences did not correlate with the strength of the enzymatic activity as assessed by labeling with the Ub-VS functional probe (Figure S2).

Upon prolonged inhibition of the RQC, stalled ribosomes may resume translation, leading to readthrough of the stall-inducing region (50). To investigate whether the inhibition of the RQC also leads to enhanced mRNA readthrough, we utilized a dual fluorescence reporter that expresses, from a single mRNA, GFP and RFP separated by a linker encoding the villin headpiece (VHP) alone (GFP-K0-RFP) or fused to a stall-inducing stretch of twenty consecutive AAA encoded Lys residues (GFP-K20-RFP) (Figure 2A). The presence of ribosomal 2A skipping sequences (P2A) on each side of the linker region allows the independent assessment of translation before and after the linker. The GFP-K0-RFP and GFP-K20-RFP reporters were cotransfected in HEK293T cells with FLAG-ev/BPLF1/UL36/UL48/Orf64 and the intensity of GFP and RFP fluorescence was measured by FACS. Based on the linear correlation of GFP and RFP fluorescence in cells expressing the GFP-K0-RFP reporter (Figure 2B, upper panels), we gated a readthrough region where the transfected cells exhibited comparable levels of green and red fluorescence, corresponding to an average RFP/GFP ratio close to 1. Upon cotransfection of the GFP-K20-RFP reporter with control FLAG-ev, robust repression of translation downstream of K20 caused the accumulation of approximately 80% of the cells into a stalling region where the RFP/GFP fluorescence ratio fell well below 1 (Figure 2B, lower left panel). Expression of vDUB rescued significant levels of RFP fluorescence (Figure 2B, lower panels). The results were highly reproducible, with 50-80% of the cells expressing the vDUBs found in the readthrough quadrant (Figure 2C). Notably, despite the high frequency of RFP rescue, the intensity of RFP fluorescence remained lower in GFP-K20-RFP/vDUB transfected compared to GFP-K0-RFP/vDUB transfected cells (compare Figure 2B, upper and lower panels). A likely explanation for this finding is that, following persistent ribosome stalling due to inhibition of the RQC, translation may restart in different frames (50), resulting in the decreased expression of fluorescent RFP species.

**Figure 2.**
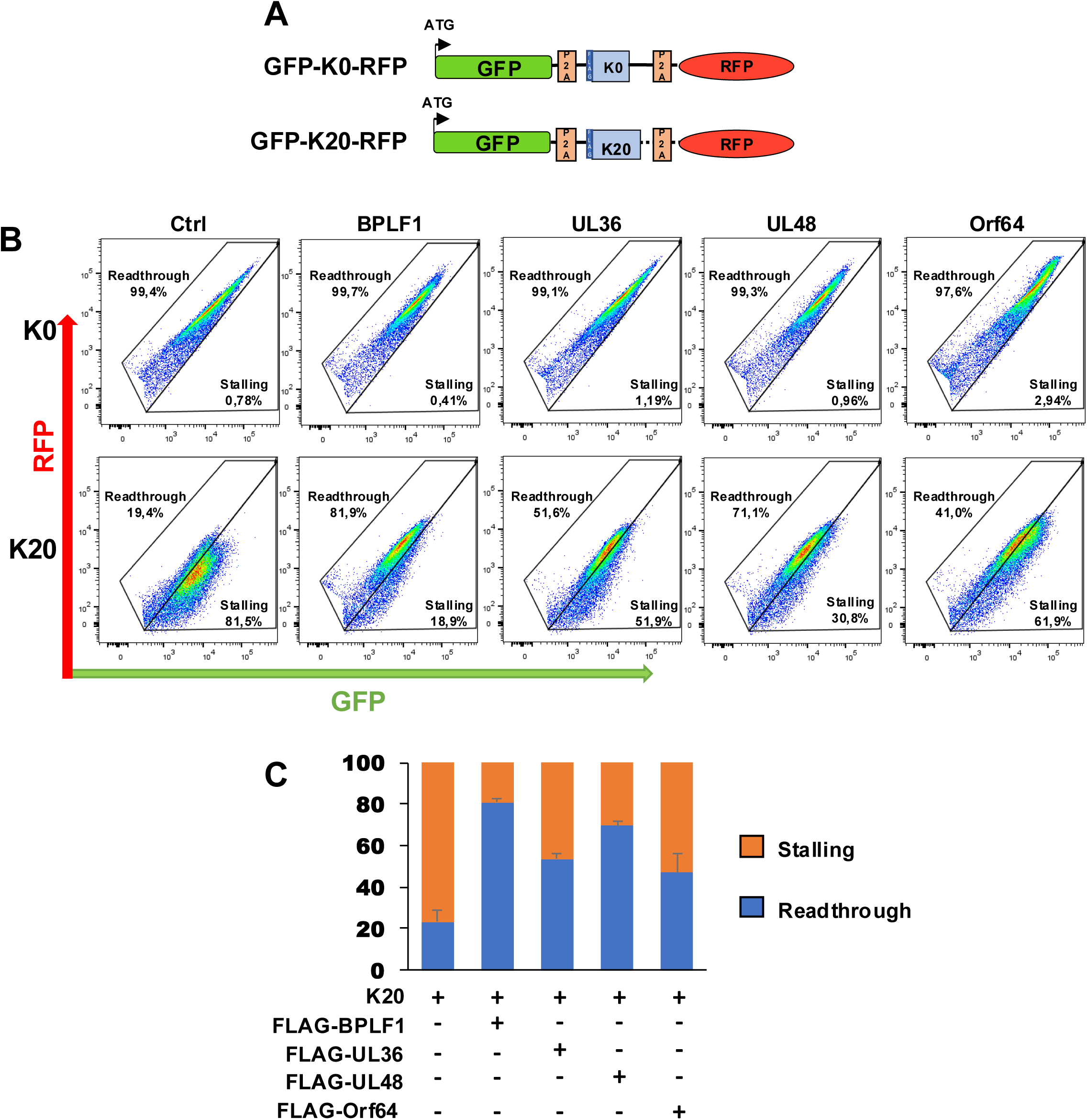
Inhibition of the RQC promotes the readthrough of stall-inducing mRNAs. **(A)** Schematic illustration of the translation-readthrough dual fluorescence reporter. The reporter expresses, from a single mRNA, the GFP and RFP separated by linker encoding a FLAG-tagged villin headpiece (VHP) alone (K0) or fused to a stall-inducing stretch of 20 consecutive lysins residues (K20). The linker regions are flanked by Ribosomal 2 A skipping sequences (P2A), which allows independent assessment of translation before and after the linker. (**B**) vDUBs promote the readthrough of stall-inducing mRNAs. FLAG-ev/vDUBs transfected HEK293T cells were co-transfected with either the K0 (upper panels) or the K20 reporter (lower panels) before GFP and RFP fluorescence analysis by FACS. After excluding non-fluorescent and dead cells, a readthrough region was defined in the plots of K0 transfected cells by gating cells exhibiting a linear correlation between GFP and RFP fluorescence. Cells falling in the stalling region exhibited decreased RFP:GFP fluorescence ratios. FACS plots from one representative experiment out of two are shown in the figure. (**C**) Mean ± SE of the percentage of transfected cells falling in the readthrough and stalling quadrants in two independent experiments.

Having established that the vDUBs share the capacity to inhibit the RQC, we then used the ER-K20 reporter to ask whether the ER-RQC is similarly affected. To this end, HEK293T cells were cotransfected with the FLAG-ev/BPLF1/UL36/UL48/Orf64 plasmids and the ER-K20 reporter (Figure 3A), and the abundance of the translated products was assessed in immunoblots probed with the GFP antibody. As controls for lysosome and proteasome-dependent degradation, FLAG-ev cotransfected cells were treated overnight with Bafilomycin A1 (BafA1) or Carfilzomib, respectively. Two faint GFP bands of approximately 50 and 48 kDa were detected in FLAG-ev transfected HEK293T cells (Figure 3B). The lower band corresponds in size to a precursor peptide arrested at the poly(A) sequence that is non-glycosylated and contains the signal peptide. This, together with the resistance to Endo-H treatment, indicates that the peptide is either released in the cytosol or accumulates at the cytosolic face of the ER. Based on sensitivity to Endo-H treatment and migration shift induced by the treatment, the upper band corresponds to an ER-localized glycosylated peptide that lacks the signal peptide (Figure 3B, red arrow). In line with the notion that luminal ER-RQC substrates are targeted for lysosome-dependent degradation, the 50 kDa species accumulated in cells treated with BafA1, whereas both Carfilzomib and BafA1 promoted the accumulation of the non-glycosylated precursor peptide. The stabilization of the non-glycosylated precursor peptide by BafA1 suggests that the fraction of the peptide that remains associated with the ER membrane, possibly trapped in the translocon, is subjected to lysosome-dependent degradation. Expression of FLAG-BPLF1, FLAG-UL48, and FLAG-Orf64 rescued the precursor and glycosylated peptides, confirming that the vDUBs inhibit both RQC and ER-RQC. In contrast, the effect of FLAG-UL36 was non-significant (Figures 3B and 3C), suggesting that the HSV1-encoded vDUB may not regulate ER-associated ribosomal stress responses.

**Figure 3.**
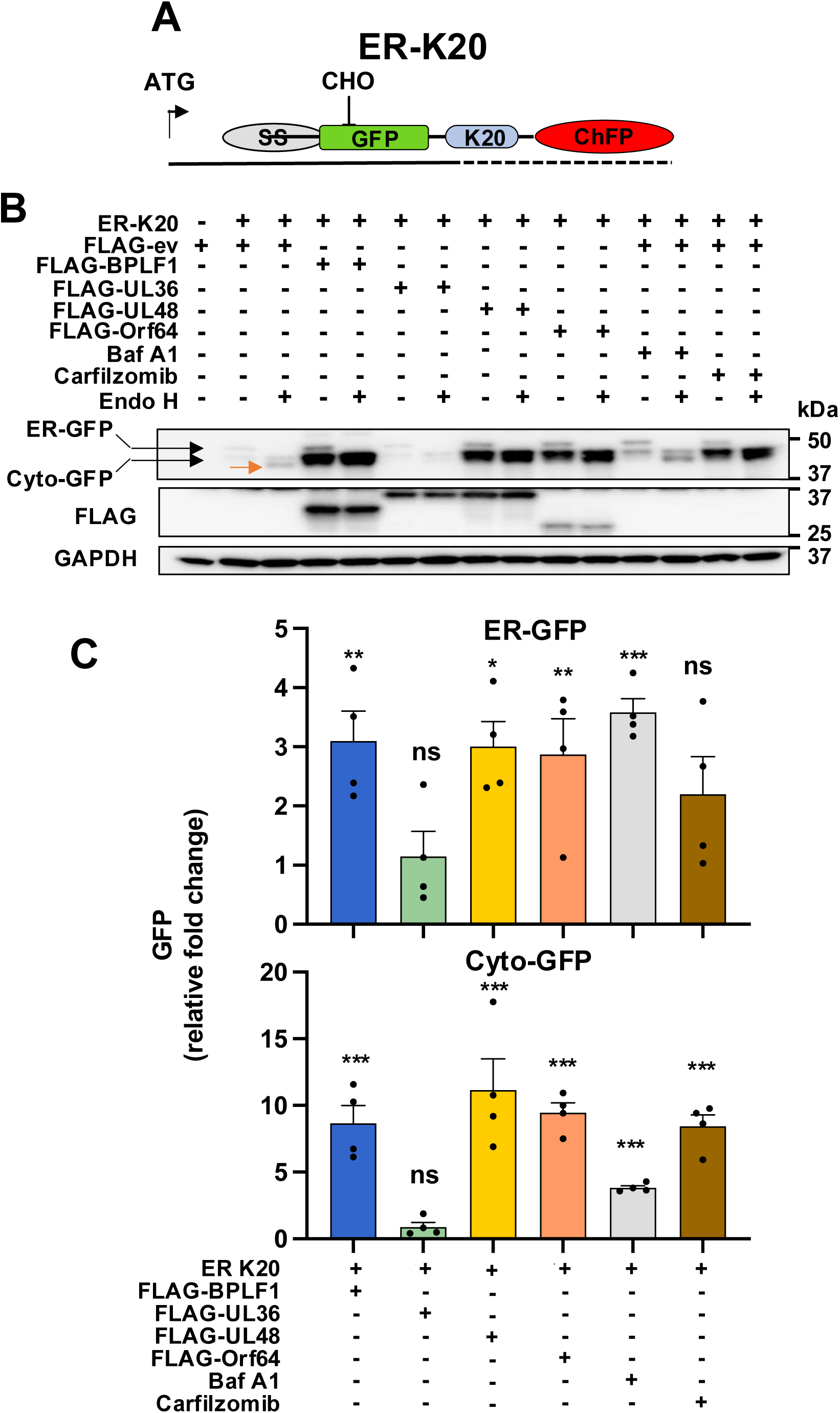
Herpesvirus deconjugase inhibit the ER-RQC. (**A**) Schematic illustration of the ER-RQC reporter (ER-K20). The reporter expresses in-frame an N-terminal ER-targeting signal sequence followed by an N-glycosylation site, GFP, and a stretch of Lys residues encoded by AAA codons (K20) and RFP. Stalling of the ribosome at the poly(A) sequence traps the nascent ER-inserted polypeptide in the translocon. **(B)** HEK293T cotransfected with the ER-K20 reporter and FLAG-ev/vDUBs were cultured for 24 h. Aliquots of FLAG-ev/ER-K20 transfected cells were treated with 100 nM Bafilomycin A1 (Baf A1) or 100 nM Carfilzomib overnight before harvesting for inhibition of lysosome- or proteasome-dependent degradation, respectively. Bands stabilized by lysosome or proteasome inhibition were indicated as ER-GFP or Cyto-GFP, respectively. Endo H was used to verify the glycosylated substrate. Immunoblots from one representative experiment out of five are shown in the figure. (**C**) Densitometry quantification of the ER-GFP or Cyto-GFP bands. Fold increase was calculated as the ratio between the band’s intensity in FLAG-vDUBs transfected cells versus FLAG-ev transfected cells after normalized to loading control. The mean± SE fold change in five independent experiments is shown. Statistical analysis was performed using an unpaired two-tailed Student’s t-test.

### HSV1-UL36 fails to inhibit ribosome UFMylation and ER-phagy

UFMylation of the 60S subunit on RPL26 is a distinctive feature of ribosomes that stall at the ER and was shown to be required to activate the ER-RQC (34). To investigate whether the vDUB regulates RPL26 UFMylation, HEK293T cells were cotransfected with S-tagged RPL26 and FLAG-ev/BPLF1/UL36/UL48/Orf64 followed by RQC activation by treatment with 50 ng/ml ANS for 1 h. The cells were then lysed in a denaturing buffer containing 20 mM NEM and 10 mM iodoacetamide to inhibit the activity of Ub/UbL deconjugases, and S-tag immunoprecipitates were analyzed by immunoblots. Three bands of sizes corresponding to RPL26 conjugated to one, two, and three UFM1 moieties were readily detected by the UFM1-specific antibody in the S-tag immunoprecipitates of ANS-treated FLAG-ev transfected cells. The UFMylation bands were virtually absent in the lysates of cells expressing FLAG-BPLF1, FLAG-UL48, or FLAG-Orf64, with a corresponding increase in the intensity of the unmodified RPL26-S band. In contrast, FLAG-UL36 had only a marginal effect, suggesting that this vDUB may lack the capacity to interfere with ER-associated UFMylation events (Figures 4A and 4B). Similar results were obtained when the UFMylation of endogenous RPL26 was assayed in ANS-treated vDUB transfected cells (Figure 4C), although the effect of the vDUBs was less prominent (Figure 4D), possibly due to the presence of a significant proportion of non-transfected cells.

**Figure 4.**
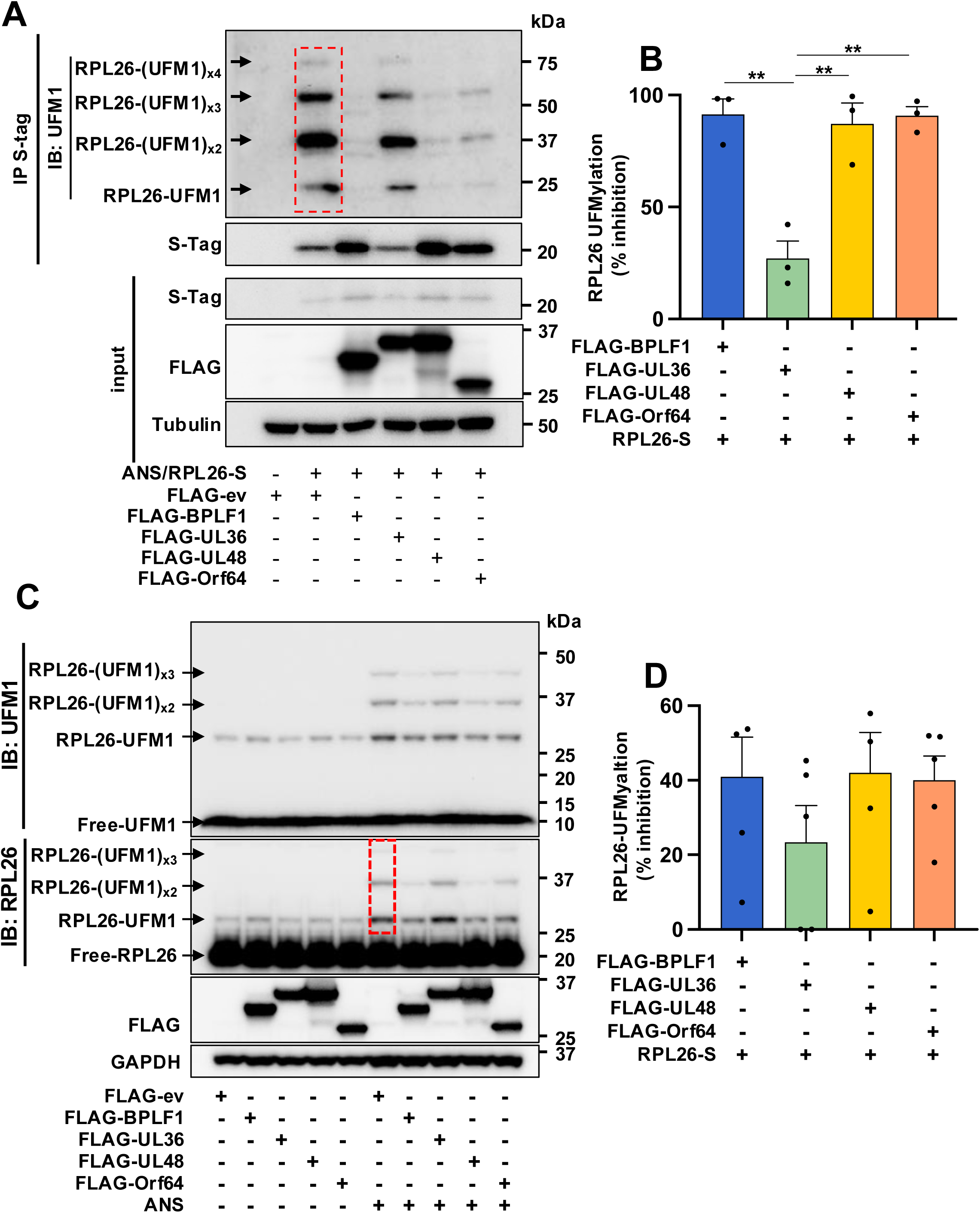
The HSV1 deconjugase does not affect the UFMylation of stalled ribosomes. (**A**) BPLF1 inhibits the UFMylation of RPL26 in ANS-treated cells. HeLa cells transfected with RPL26-S Tag alone or together with FLAG-ev/vDUBs as indicated were treated with 50 ng/ml ANS for 30 min, followed by lysis under denaturing conditions to destroy non-covalent interactions and immunoprecipitation with anti-S-coupled beads. Arrows indicate mono- di- tri- and tetra-UFMylated RPL26 detected by UFM1 antibody. Immunoblots from one representative experiment out of three are shown in the figure. (**B**) Densitometry quantification of the UFMylated bands. The area included in the densitometry scan is indicated by a red dotted box in Figure 4A. Percentage inhibition was calculated relarive to FLAG-ev transfected cells. The significance of the differences between the effect of HSV1-UL36 versus the homologs was calculated by unpaired two-tailed Student’s t-test. (**C**) The vDUBs inhibit the UFMylation of endogenous RPL26 induced by ANS treatment. HEK293T cells transfected for 24 h with FLAG-ev/vDUBs were treated with 50 ng/ml ANS for 30 min, followed by lysis in a buffer containing NEM to inhibit DUB activity. High molecular weight species corresponding to the size of mono- di- and tri-UFMylated RPL26 (arrows indicated) were detected in immunoblots probed with antibodies specific for UFM1 or RPL26. Blots from one representative experiment out of four are shown in the figure. (**D**) Densitometry quantification of the UFMylated species. The area included in the densitometry scan is indicated by a red dotted box in Figure 4C. The intensity was normalized to the intensity of the RPL26 band in short exposure of the same blots. The mean ± SE of four independent experiments is shown.

The UFMylation of ribosomes and ER-membrane proteins was shown to regulate ER-phagy, which participates in the clearance of ER-RQC substrates (51). To test whether the effect of the vDUBs on ribosome UFMylation impacts ER-phagy, we used the previously described subline of HCT116 expressing a Dox-regulated ER-phagy dual fluorescence reporter constructed by in-frame fusion of the coding sequences of the RAMP4 subunit of the ER translocon, enhanced GFP (eGFP), and mCherry Fluorescent Protein (ChFP) (46) (Figures 5A). Upon ER insertion of the RAMP4 domain, GFP and ChFP face the cytosol and emit equal fluorescence, whereas, due to the selective loss of GFP fluorescence at low pH, ER-loaded autophagolysosomes appear as distinct red fluorescent dots (Figure S3). To assess the effect of the vDUB on this process, the reporter cell line was transfected with plasmids expressing FLAG-BPLF1/UL36/UL48/Orf64 and cultured for 24 h in the presence of Dox before starvation overnight in EBSS medium followed by visualization of autophagosomes by confocal microscopy. Analysis of red cargo-loaded phagolysosomes in transfected and non-transfected cells from the same slide resulted in the ready detection of red dots in the non-transfected cells. However, only a diffuse yellow fluorescence was observed in most cells expressing FLAG-BPLF1, FLAG-UL48, or FLAG-Orf64. In contrast, red dots were clearly visible in cells expressing FLAG-UL36 (Figure 5B). Quantification of the number of red dots in positive and negative cells from the same transfection revealed highly significant suppression of ER-phagy in cells expressing FLAG-BPLF1/UL48/Orf64 (Figure 5C) whereas, although somewhat decreased in numbers, dots were still detected in that majority of FLAG-UL36 expressing cells.

**Figure 5.**
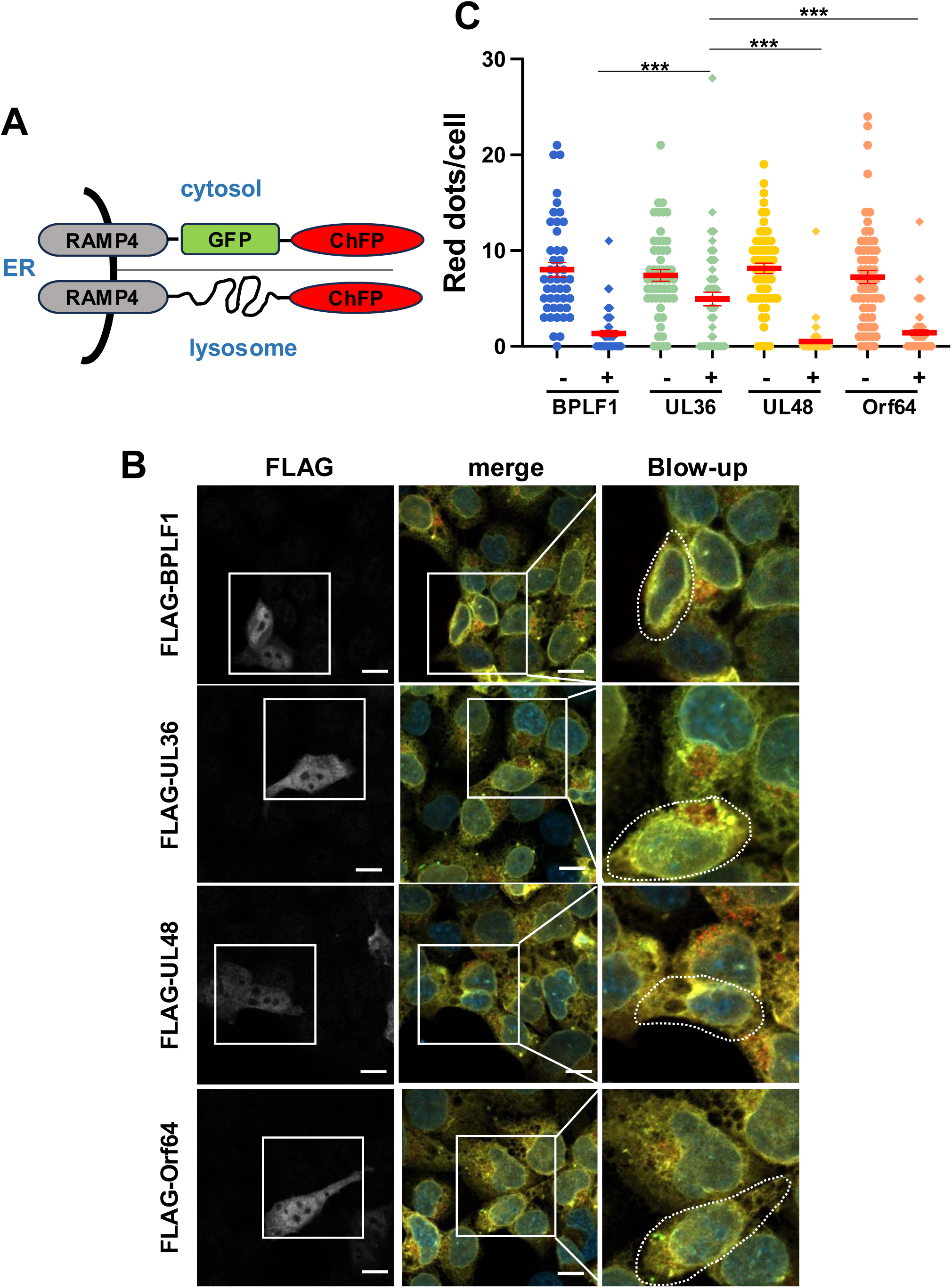
The HSV1 deconjugase does not inhibit ER-phagy. (**A**) Schematic illustration of the ER-Autophagy Tandem Reporter (EATR). The reporter expresses in-frame the coding sequence of the RAMP4 subunit of the ER translocon complex followed by the coding sequences of eGFP and ChFP. Upon ER insertion of the RAMP4 domain, eGFP and ChFP face the cytosol and emit equal fluorescence, whereas, due to the selective loss of eGFP fluorescence at low pH, ER-loaded autophagosomes appear as distinct red fluorescent dots. (**B**) Representative confocal images illustrating the failure to accumulate red fluorescent dots in cells expressing vDUBs starved overnight, respectively. Stable HCT116-EATR cells were transfected with plasmids expressing FLAG-vDUBs and then starved overnight in EBSS medium before visualizing the formation of ER-loaded autophagosomes by confocal microscopy. (**C**) Quantification of the number of red fluorescent dots in vDUBs positive and negative cells from the same transfection experiments. The cumulative data from two independent experiments where approximately 50 vDUB positive and negative cells scored from the same slide are shown. Significance was calculated using an unpaired two-tailed Student t-test.

## Discussion

The site-specific post-translational modification of ribosomal proteins by the covalent attachment of ubiquitin and ubiquitin-like polypeptides plays a pivotal role in the regulation of the RQC and the ribosome and integrated stress responses that are activated by the hindrance of translation initiation, elongation or termination (52). The conservation of these stress-induced modifications across species points to the involvement of individual events in the regulation of different steps in the translation cycle. Indeed, while the coordinated ubiquitination of RPS10 and RPS20, and RPL26 UFMylation was shown to be critical for the activation of RQC or ER-RQC during translation elongation, a different type of RQC, characterized by the coordinated ubiquitination of RPS3 and RPS2, is activated by events that hamper translation initiation (iRQC) (53), or activate the ISR (15, 16). The relationship between these different types of translational stress responses, their downstream effects, and physiological functions are still poorly understood, but it is generally acknowledged that while the triggering of RQC or ER-RQC promotes the degradation of aberrant translation products and the recycling of ribosome subunits, the coordinated ubiquitination of RPS3 and RPS2 is accompanied by disposal of 40S subunit via proteasome-dependent degradation (16).

Here, we have found that the deconjugases encoded in the N-terminal domain of the large tegument protein of the human pathogenic herpesviruses EBV, HCMV, and KSHV share the capacity to inhibit various branches of the RQC by impairing the ubiquitination of RPS10, RPS20 and RPS3 and UFMylation of RPL26 while the HSV1 encoded vDUB selectively failed to prevent RPL26 UFMylation and the activation of ER-phagy. The coordinated inhibition of different types of RQC responses may offer several advantages to the viruses. One important outcome is likely to be the boosting of proficient translation of viral mRNA containing a variety of translational hindrances, as illustrated by the capacity of the vDUBs to promote the readthrough of stall-inducing mRNAs. Conceivably, the stabilization of paused/stalled ribosomes due to the combined effect of failure to initiate the disassembly of stalled ribosomes via the recruitment of ASC-1 to ubiquitinated RPS10/RPS20 and stabilization of the 40S subunit through inhibition of RPS2/RPS3 ubiquitination-triggered degradation is critical for enabling the restart of translation after prolonged pause, which may be particularly important for the translation of viral mRNAs. In line with this possibility, we have previously shown that the enzymatic activity of the EBV encoded BPLF1 is required for the strong promoting effect of the vDUB on the translation of the viral genome maintenance protein EBNA1 whose mRNA contains a long G-C rich domain that forms G-quadruples and inhibits translation elongation (54). Notably, a similar long repeat is found in the KSHV-encoded genome maintenance protein LANA1 (55) and shorter G-quadruplex forming repeats are present in the protein-encoding mRNAs of several herpesviruses including HCMV and HSV1 (56), pointing to a conserved function of the vDUBs in promoting the translation of viral proteins.

We have previously reported that vDUB encoded by EBV, HCMV, and KSHV activate the ISR via the ZAKα and GCN2 mediated phosphorylation of eIF2α while the HSV1 DUB failed to do so (45). Together with the iRQC inhibition underlined by the repression of RPS3 ubiquitination identified in this work, the activation of the ISR is likely to play an important role in the reprogramming of translation towards mRNA with alternative start sites such as 5’ untranslated open reading frames (uORFs) and internal ribosomal entry sites (IRES) that are frequently found in viral mRNAs (42, 57) and in the mRNAs in cellular genes with growth-promoting and anti-apoptotic functions (42, 43, 58). In support of this scenario, the effect of BPLF1 on the translation of the EBNA1 mRNA expressed from a lytic promoter that contains both uORFs and IRES signatures (57, 59) was selectively blocked by the inhibition of GCN2 but not by other ISR kinases (45). Interestingly, while we now find that the UL36 inhibits the ubiquitination of RPS3, albeit somewhat less efficiently than the homologs, we have previously shown that the HSV1 encoded DUB does not activate the ISR, as assessed by the failure to promote the upregulation of ATF4 (45). Thus, activation of the ISR may not be required to trigger RPS3 ubiquitination in herpesvirus-infected cells, which may be dependent on events occurring during the translation cycle. These may include a prolonged stalling of the 43S preinitiation complex during the scanning of viral mRNAs or collision of the scanning complex with stalled 80S ribosome as suggested for the effect of chemical compounds that trigger the iRQC (53).

We have found that in addition to inhibiting RQC and iRQC, the vDUBs encoded by EBV, HCMV, and KSHV share the capacity to inhibit the ER-RQC as illustrated by the inhibition of RPL26 UFMylation, rescue of a model ER-RQC substrate from both proteasome and lysosome-mediate degradation and of inhibition of ER-phagy. The inhibition of this branch of the RQC may provide an additional advantage to the viruses by enhancing the translation of viral glycoproteins that play a critical role in the assembly of infectious virus particles. Although its involvement in both ER-RQC and ER-phagy is firmly established (34, 51, 60), the mechanism by which RPL26 UFMylation promotes these events is not fully understood. It has been proposed that UFMylation may be required to relax the ribosome-translocon junction, exposing the trapped polypeptide to degradation by the cytosolic RQC (34). This scenario aligns with the finding that RPL26 UFMylation plays a physiological role in promoting the release of ribosomes from the ER upon translation termination (61, 62). The triggering event is also unclear since RPL26 UFMylation is not affected by ablation of ZNF598, the prime reader of ribosome stalling and collision (46). We have previously shown that BPLF1 acts by reversing constitutive or induced ubiquitination events required for RPL26 UFMylation (46), which correlated with strongly decreased ubiquitination of ribosome and associated proteins that comigrate in sucrose gradient. In this context, the failure of HSV1 UL36 to regulate the UFMylation of RPL26, stabilize ER-RQC substrates, and inhibit the activation of ER-phagy was surprising, given the comparable enzymatic activity of viral enzymes. The discrepancy is likely explained by differences in the substrate repertoire of the HSV1 encoded enzymes, as also suggested by our previous finding that, different from the homologs, UL36 fails to regulate IFN responses via interaction with 14-3-3 proteins (63) and does not inhibit macroautophagy via deubiquitination of the SQSTM/p62 receptor (64). This possibility is supported by modeling of the DUB domain of the tegument proteins, which revealed a striking correlation between the divergent substrate repertoire of UL36 and the length and charge distribution of the solvent-exposed residues in helix-2 (65) that was shown to be critical for the substrate interaction of BPLF1 (66).

During productive infections, viruses are strictly dependent on the cellular mRNA translation machinery for the synthesis of proteins required for virus replication and the assembly of new virus particles. While a variety of viral strategies for diverting the cellular translation apparatus toward the mass production of comparatively small viral proteomes have been intensely studied, how viruses deal with the potentially antiviral effects of quality control systems that ensure the fidelity of ribosome translation and the disposal of aberrant translation products has not been systematically investigated. Our findings provide a first insight into this issue in the context of infection by human pathogenic herpesviruses by pointing to an important contribution of the virus-encoded vDUBs in the regulation of ubiquitination events that control different types of RQC responses. It is tempting to speculate that the different behavior of the HVS1 encoded DUB, and in particular, the failure to counteract ER-associate translation quality control and the ISR responses may reflect the different host cell range and substantial differences in the length and regulation of the replicative cycle.

## Data Availability

All data that support the findings of this study are contained within the manuscript and are available on reasonable request.

## Open Access

Open access for this article was enabled by the participation of the Karolinska Institute in an all-inclusive Read & Publish pilot with Portland Press and the Biochemical Society.

## CRediT Author Contribution

Jianagnan Liu: Conceptualization, formal analysis, investigation, visualization, and original draft writing. Carlos Ayala-Torres: investigation, data analysis. Maria G. Masucci: administration, data analysis, writing, review, and editing.

## Acknowledgments

The technical input of the master students Francisco Morais Estevez and Maitri Paul is gratefully acknowledged. The Masucci lab is a member of the COST network ProteoCure.

## Abbreviations

EBV, Epstein–Barr virus; BPLF1, BamH1 fragment left open reading frame-1; HSV1, herpes simplex virus-1; UL36, unique long 36; HCMV, human cytomegalovirus, UL46, unique long 48; KSHV, Kaposi sarcoma herpesvirus; Orf64, open reding frame 64; DUB, deubiquitinating enzyme; USP, ubiquitin specific protease; RQC, ribosome quality control, iRQC, initiation RQC; ER, endoplasmic reticulum, ER-RQC, ER-associated RQC; ISR, integrated stress response; RSS, ribosomal stress response; UFM1, ubiquitin family modifier-1; NEMF, nuclear export mediator factor; VCP/p97, valosin containing protein/p97

