## supplementary table s1 and Figure S1,S2,S3 for "Differential regulation of translational stress responses by herpesvirus ubiquitin deconjugases"

**Supplementary information**

**Supplementary Tables**

**Table S1. Reagents used in this paper**

| **Reagent** | | **Source** | **Identifier** |
|---|---|---|---|
| **Antibodies** | **Working dilutions** | | |
| Mouse monoclonal Anti-β-actin clone AC-15 | 1:5000 | Sigma-Aldrich | Cat# A5441, RRID: AB_476744 |
| Mouse monoclonal Anti-GAPDH | 1:10000 | Millipore | Cat#CB1001 |
| Mouse monoclonal anti-FLAG | 1:10000 | Sigma-Aldrich | Cat# F3165, RRID: AB_259529) |
| Rabbit polyclone anti-FLAG | 1:5000 | Sigma-Aldrich | Cat#F7425, RRID: AB_439687 |
| Mouse monoclonal anti-S Tag, | 1:1000 | Millipore | Cat#71549-3 RRID: AB_11210600 |
| Mouse monoclonal anti-HA Tag | 1:1000 | Sigma-Aldrich | Cat# H9658 RRID:AB_260092 |
| S-Protein Agarose | | Novagen | Cat#69704 |
| Goat Anti-Rabbit IgG (H+L) Antibody, Alexa Fluor 647 Conjugated | 1:1000 | Thermo Fisher Scientific | Cat#A21245 RRID: AB_141775 |
| Rabbit Monoclonal anti-UFM1 | 1:2000 | Abcam | Cat# ab109305, RRID: AB_10864675 |
| Rabbit Polyclonal anti-RPL26 | 1:3000 | Abcam | Cat# ab59567, RRID: AB_945306 |
| Rabbit polyclonal anti-ZNF598 | 1:5000 | Abcam | AB241092 |
| Rabbit monoclonal anti-RPS10 | 1:2000 | Abcam | Cat# ab151550 RRID:AB_2714147 |
| Rabbit monoclonal anti-RPS20 | 1:3000 | Abcam | Cat# ab133776 RRID:AB_2714148 |
| Rabbit monoclonal anti-RPS3 | 1:2000 | Abcam | Cat# ab128995 RRID:AB_11145466 |
| Mouse monoclonal anti-GFP | 1:2000 | Santa Cruz | Cat# sc-9996 RRID:AB_627695 |
| Mouse monoclonal anti-ubiquitin | 1:1000 | Santa Cruz | Cat#sc-8017 RRID:AB_628423 |
| **Chemicals, peptides, and recombinant proteins** | | | |
| IGEPAL CA-630 | Sigma-Aldrich | | I3021; CAS: 9002-93-1 |
| Ciprofloxacin | Sigma-Aldrich | | Cat#17850; CAS: 85721-33-1 |
| $MgCl_2$ | Merck | | Cat# M1028 |
| Sodium dodecyl sulphate | Sigma-Aldrich | | L3771; CAS:151-21-3 |

| | | |
|---|---|---|
| Sodium deoxycholate monohydrate | Sigma-Aldrich | D5670; CAS:145224-92-6 |
| Triton X-100 | Sigma-Aldrich | T9284; CAS:9002-93-1 |
| Bovine serum albumin | Sigma-Aldrich | A7906; CAS:9048-46-8 |
| Tween-20 | Sigma-Aldrich | P9416; CAS: 9005-64-5 |
| PMSF | Sigma-Aldrich | P7626; CAS: 329-98-6 |
| Trizma base | Sigma-Aldrich | 93349; CAS:77-86-1 |
| Doxycycline cyclate | Sigma-Aldrich | D9891; CAS: 24390-14-5 |
| Anisomycin | Sigma-Aldrich | A5862; CAS:22862-76-6 |
| Carfilzomib | Medchem Express | Cat# 253339 |
| Bafilomycin A1 | Sigma-Aldrich | B1793; CAS:88899-55-2 |
| Complete protease inhibitors cocktail | Roche Diagnostic | Cat#04693116001 |
| DAPI | Sigma-Aldrich | Cat#9542 CAS:28718-90-3 |
| Mowiol | Calbiochem | Cat# 475904 CAS:9002-89-5 |
| Dabco(1,4-Diazabicyclo[2.2.2]octane) | Sigma-Aldrich | Cat#D2522 CAS:280-57-9 |
| HA-Ubiquitin-VS | R&D systems | Cat# U-212-025 |
| **Kits** | | |
| jetOPTIMUS DNA transfection reagent | Polyplus | Cat#101000006 |
| DC Protein Assay quantification kit | Bio-Rad | Cat#A500-0116 |
| SuperSignal$^{TM}$ WestPico PLUS Chemiluminescent Substrate | Thermo Scientific | Cat#XE356732 |
| Endo H | New England biolabs | Cat# P0702S |
| **Medium and Buffer** | | |
| DMEM | Sigma-Aldrich | Cat# D6429 |
| EBSS | Gibco | Cat# 1854705 |
| **Experimental models: Cell lines** | | |
| HEK293T | ATCC | CRL3216 |
| HeLa | ATCC | RR-B51S |
| HCT116 | ATCC | CCL-247 |
| HCT116-EATR | This paper | N/A |
| HEK293T *ZNF598* knockout | Liu et al., 2023 | N/A |
| **Plasmids** | | |
| TetOn-mCherry-eGFP-RAMP4 | gift from Jacob Corn | Addgene #109014 |
| ER-K20 | gift from Yihong Ye | Addgene # 133861 |
| K20 | This paper | N/A |
| pmGFP-P2A-K0-P2A-RFP | gift from Ramanujan Hegde | Addgene # 105686 |
| pmGFP-P2A-K(AAA)20-P2A-RFP | gift from Ramanujan Hegde | Addgene # 105688 |
| pCMV10-3xFLAG-BPLF1 | Li et al. PloS Path., 2021 | N/A |
| pCMV10-3xFLAG-UL36 | Gupta et al. Front Immunol., 2021 | N/A |

| | | |
|---|---|---|
| pCMV10-3xFLAG-UL48 | Gupta et al. Front Immunol, 2021 | N/A |
| pCMV10-3xFLAG-Orf64 | Gupta et al. Front Immunol., 2021 | N/A |
| pCDNA3.1-RPL26-S | gift from Ron R. Kopito (Stanford University, Stanford, CA, USA) | N/A |
| **Primers used for cloning K20** | | |
| Forward: GTGAGCAAGGGCGAG | Reverse: CATGGTGGCGACCGG | |

**Supplementary Figure Legends**

**Figure S1. The vDUBs inhibit the ubiquitination of endogenous RPS10 and RPS20 in ANS-treated cells.** (**A**) FLAG-ev/vDUBs transfected HEK293T cells were cultured for 24 h and then treated with 50 ng/ml ANS for 30 min before lysis in buffer containing NEM and iodoacetamide to inhibit DUB activity. Representative blots from one out of four independent experiments are shown in the figure. (**B**) Densitometry quantification of the intensity of the ubiquitinated species. The area included in the densitometry scan is indicated by a red dotted box in Figure S1. The mean ± SE % inhibition relative to ANS-treated FLAG-ev transfected cells in four independent experiments is shown.

**Figure S2**. **Assessment of enzymatic activity by Ub-VS labeling.** (**A**) HEK293T cells were transfected with the indicated FLAG-vDUB vectors. After 24 h, the cells were lysed, and equal amounts of cell lysates were incubated with 1 µM of the HA-Ub-VS functional probe at 37 °C. The reactions were stopped at the indicated times by adding NuPAGE loading buffer, and western blots were probed with the FLAG antibody. Enzymatic activity is indicated by a 10 kDa migration shift corresponding to the covalent attachment of the probe to catalytic Cys residue. (**B**) Densitometry quantification of the percentage of the Ub-VS conjugates vDUB relative to the total (labeled plus unlabeled). One representative experiment out of two is shown.

**Figure S3**. **Characterization of the ER-phagy reporter cell line**. HCT116-EATR were grown overnight in complete medium or starvation-inducing EBSS medium in the presence or absence of BafA1 (100 nM). When grown in a complete medium, the reporter exhibited a diffuse yellow signal, indicating equal expression of GFP and mCherry. Upon starvation-induced ER-Phagy, red dots corresponding to ER-loaded autophagosomes became apparent due to the quenching of GFP fluorescence caused at low pH conditions. When acidification was hindered by the presence of Bafilomycin A1, yellow dots were observed.

### Figure S1

A

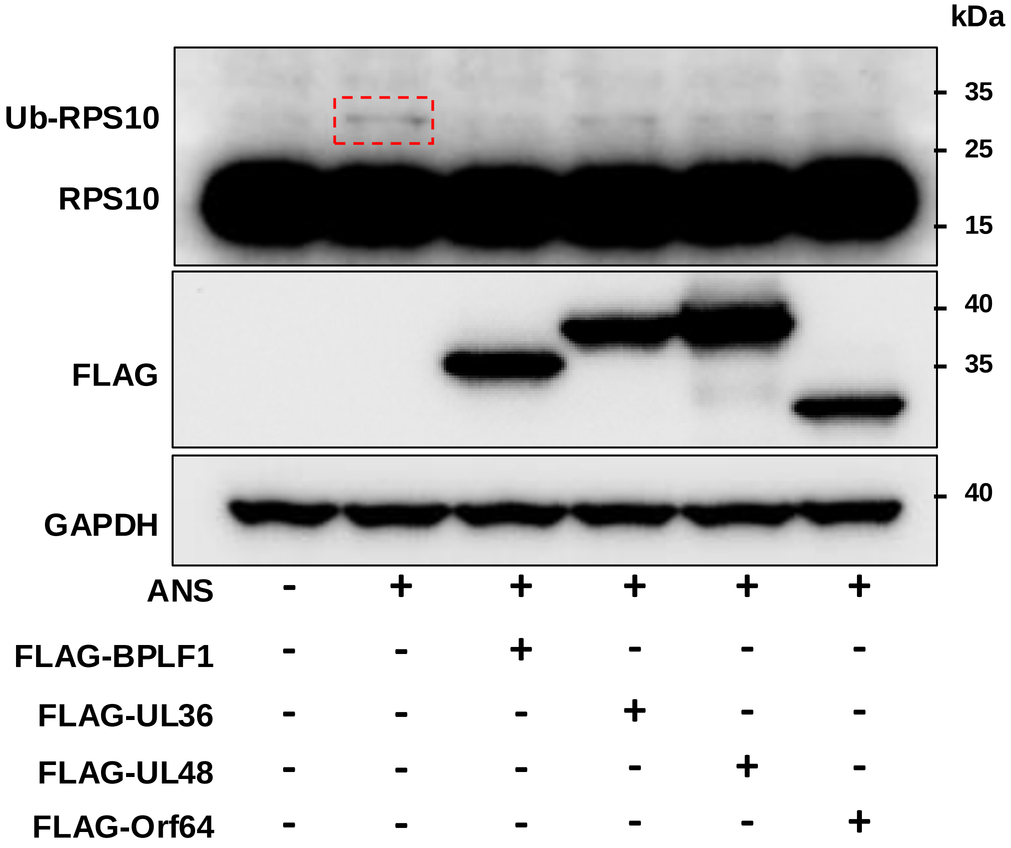
kDa
Ub-RPS10
RPS10
FLAG
GAPDH
35
25
15
40
35
40
ANS - + + + + +
FLAG-BPLF1 - - + - - -
FLAG-UL36 - - - + - -
FLAG-UL48 - - - - + -
FLAG-Orf64 - - - - - +

B

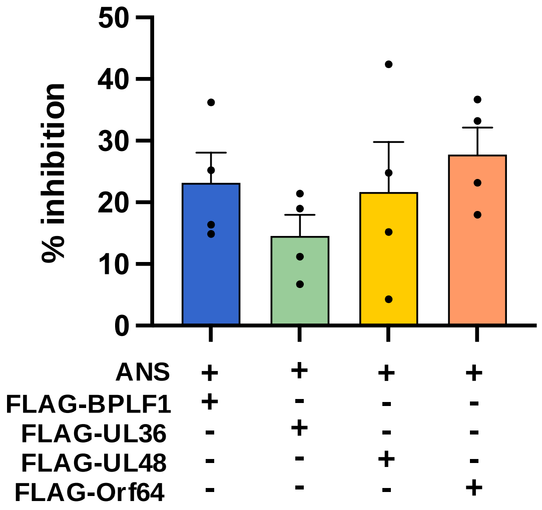
% inhibition
50
40
30
20
10
0
ANS + + + +
FLAG-BPLF1 + - - -
FLAG-UL36 - + - -
FLAG-UL48 - - + -
FLAG-Orf64 - - - +

### Figure S2

A

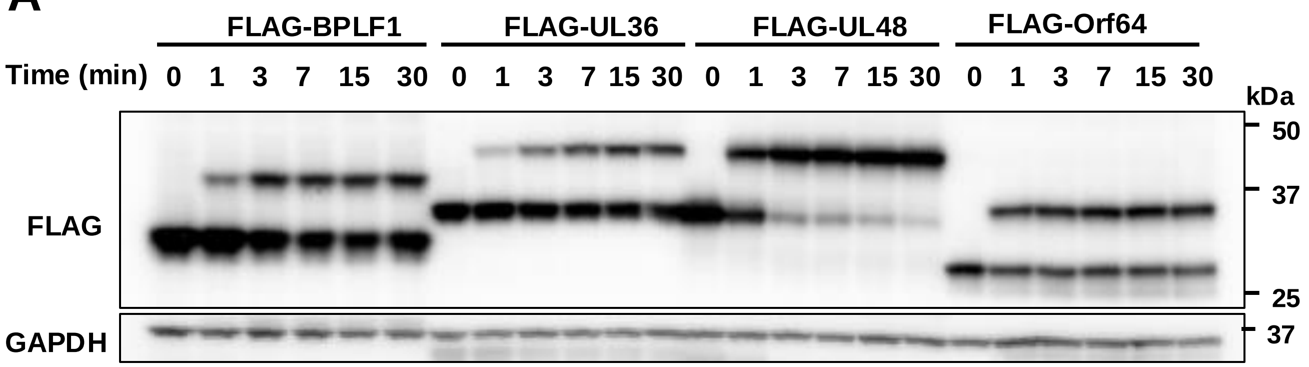
FLAG-BPLF1
FLAG-UL36
FLAG-UL48
FLAG-Orf64
Time (min) 0 1 3 7 15 30 0 1 3 7 15 30 0 1 3 7 15 30 0 1 3 7 15 30
kDa
50
37
25
37
FLAG
GAPDH

B

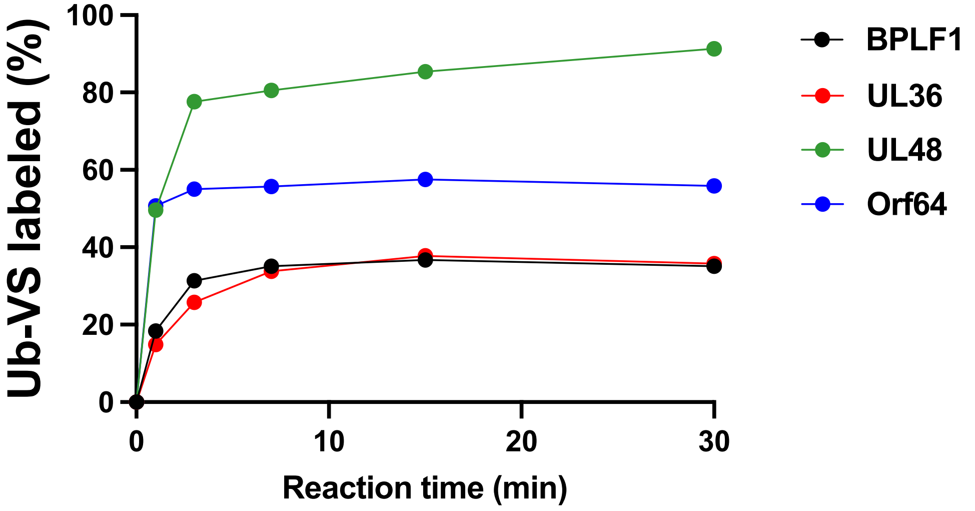
Ub-VS labeled (%)
100
80
60
40
20
0
0
10
20
30
Reaction time (min)
BPLF1
UL36
UL48
Orf64

### Figure S3

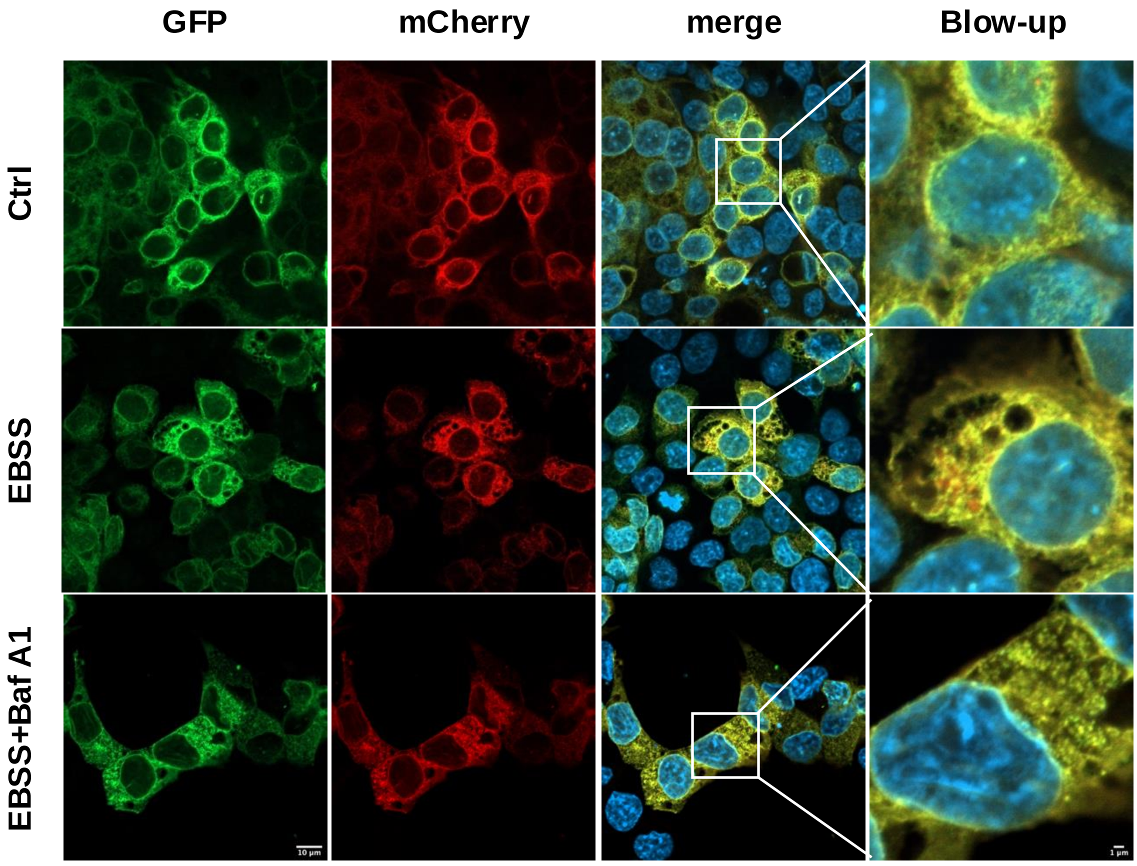
